# Elevated rates of molecular evolution genome-wide in mutualist legumes and rhizobia

**DOI:** 10.1101/2024.06.10.598267

**Authors:** L. T Harrison, J. R Stinchcombe, M. E Frederickson

## Abstract

Rates of molecular evolution vary greatly among even closely related species. Although theory predicts that antagonistic interactions between species increase rates of molecular evolution, predictions for how mutualism affects evolutionary rates are mixed. Here, we compared rates of molecular evolution between 1) mutualistic and non-mutualistic legumes, 2) an independent set of symbiotic rhizobia and their non-symbiotic close relatives, and 3) symbiotic and non-symbiotic clades within *Ensifer*, a diverse genus of bacteria with various lifestyles. We assembled transcriptomes *de novo* for 12 legume species and then calculated dN/dS ratios at orthologous genes in all species to determine if genes in mutualistic plants evolve faster or slower than in their non-mutualistic relatives. We also calculated dN/dS ratios in symbiosis genes known to be important for nodulation with rhizobia. We found that mutualists have higher rates of molecular evolution genome-wide compared to non-mutualist legumes. We next calculated dN/dS ratios in 14 bacteria species across the proteobacteria phylogeny that differ in whether they associate mutualistically with plants, using previously published data. We found that in most pairs, symbiotic rhizobia show higher dN/dS values compared to their non-symbiotic relatives. Finally, within a bacterial genus with many well-characterized mutualist species (*Ensifer*), we calculated dN/dS ratios in symbiotic and non-symbiotic clades and found that symbiotic lineages have higher rates of molecular evolution genome-wide, but not at genes on the symbiotic plasmid pSymB. Our results suggest that although mutualism between legumes and rhizobia is associated with elevated rates of molecular evolution genome-wide, symbiosis genes may be evolutionarily stagnant.

## Introduction

A fundamental question in evolutionary biology is how species interactions contribute to variation in rates of molecular evolution (Woolfit and Bromham 2003; Bromham 2009). According to the Red Queen hypothesis, species interacting antagonistically will have higher rates of molecular evolution because they are under constant pressure to evolve new defenses against their enemies, or new counter-adaptations that overcome their victims’ defenses (Stahl et al. 1999; Brockhurst et al. 2014; Delaye et al. 2018). It is less clear how mutualism might impact molecular evolution. On the one hand, mutualists might have higher rates of molecular evolution than non-mutualistic species because they have to adapt to both a changing environment and a changing partner (Lutzoni and Pagel 1997; Rubin and Moreau 2016). On the other hand, some theory suggests that the more slowly evolving partner in a mutualism reaps the greatest rewards (the so-called Red King hypothesis, Bergstrom and Lachmann 2003). Slower rates of evolution are also expected if mutualist partners reach evolutionary stasis (Hembry et al. 2014; Barker et al. 2017), when selection maintains interacting species near trait-matched fitness optima with little further phenotypic change (Nuismer et al. 2013). In this scenario, stabilizing selection on both hosts and rhizobia would result in fewer nucleotide substitutions in the genome, particularly at symbiosis genes (Epstein et al. 2022). In addition, if most genetic variation in mutualist quality is due to mutation-selection balance, a signature of purifying selection would be expected, at least at symbiosis genes (Heath and Stinchcombe 2014). Although population genetic methods can identify neutral and selective pressures on traits (Tiffin and Ross-Ibarra 2014; O’Brien et al. 2021), it is challenging to distinguish between stabilizing and purifying selection with population genetic data (Charlesworth 2013). Here, we compare rates of molecular evolution between 1) mutualistic and non-mutualistic legumes, 2) taxonomically diverse symbiotic and non-symbiotic rhizobia, and 3) symbiotic and non-symbiotic clades within a single well-studied rhizobia genus (*Ensifer)* to determine whether the legume-rhizobium symbiosis involves rapid or slow DNA sequence evolution.

The facultative legume-rhizobium mutualism, in which leguminous plants exchange carbon for fixed nitrogen provided by rhizobial partners, is an excellent system to test how mutualism influences rates of molecular evolution. Rhizobia occupy specialized root structures on legumes called nodules (van Rhijn and Vanderleyden 1995), and this symbiosis is generally mutualistic (Friesen 2012) especially in low-nitrogen environments. The legume family (Fabaceae/Leguminosae) is large, including around 19500 species (Azani et al. 2017), but not all these species form nodules with rhizobia. Although some work suggests that nodulation in plants has evolved multiple times after a single predisposition event (Doyle 2011; Werner et al. 2014), recent phylogenomic analyses support the hypothesis that nodulation has a single evolutionary origin, followed by multiple losses of the trait across the clade (Griesmann et al. 2018, Parshuram et al. 2023).

There is also high variation in nodulation capabilities among bacteria. Rhizobia are horizontally transmitted symbionts that are taken up from the soil by new legume hosts each generation, meaning rhizobial lineages alternate between being plant-associated and free-living in soil. Bacterial strains that have nodulation genes (*nod* genes) produce Nod factors that are important for initiating nodule formation on plant roots. However, not all rhizobia with *nod* genes can form nodules on all legume species; legumes have Nod factor receptors that must recognize compatible Nod factors for a successful symbiosis to occur (Wang et al. 2018). The development of nodules and nitrogen fixation are complex processes, requiring many genes (*nif*, *fix*, etc.) that are often found together on a mobile genetic element such as a plasmid (Batstone 2022). Rhizobia can exchange these symbiosis genes or plasmids, and thus symbiotic ability, through horizontal gene transfer (Epstein and Tiffin 2021; Rahimlou et al. 2021).

Many genetic changes that accompany transitions to mutualism have been identified in endosymbiotic bacteria. For example, extremely tiny genomes are a common feature of obligately endosymbiotic bacteria that are vertically transferred to new hosts (McCutcheon and Moran 2012). These bacteria rely on their host for many functions and thus many genes are lost in their own genome (Wernegreen 2002). Endosymbiotic bacteria also experience bottlenecks each time they are passed down to a new host (Woolfit and Bromham 2003). The reduction in population size leads to a decrease in genetic variation in the new population and a greater chance that variants are fixed or lost due to this random sampling, leading to higher rates of nucleotide substitution. Although rhizobia are also endosymbiotic within plant cells, they have a free-living stage, are horizontally transmitted, and may gain nodulation abilities through horizontal gene transfer, making it less clear how mutualism will impact genome and molecular evolution. Associating with diverse plant species and spending some time in the soil without a host might weaken host selection on the rhizobia genome (Sachs et al. 2011). Many of the genomic signatures of coevolution might be observed only in symbiosis genes, if these genes are commonly horizontally transferred into the genomes of non-symbiotic lineages (Epstein et al. 2022).

In plants, we might expect that many adaptive mutations would be necessary for the evolution of nodulation, resulting in signals of positive selection in mutualist lineages. If nodulation evolved only once near the base of the legume tree (Griesmann et al. 2018), strong positive selection may have occurred in response to mutualists in the past, but may no longer be detectable with population genetic methods. Previous work has shown that the evolution of polyploidy in legumes likely pre-dated symbiosis and may have facilitated the evolution of nodulation (Parshuram et al. 2023), suggesting that nodulation is not easily gained in multiple lineages. Nonetheless, around 9% of legumes do not form nodules (Simonsen et al. 2017), with a phylogenetic distribution that suggests multiple losses of this trait. When nodulation is lost, we might expect relaxed selection on genes that were formerly important for symbiosis with rhizobia and thus higher rates of molecular evolution at symbiosis genes in non-mutualistic lineages. In addition, mutualism is expected to increase population sizes by allowing organisms to thrive despite enemies, abiotic stress, or nutrient limitation (Afkhami et al. 2014; Weber and Agrawal 2014). Consequently, purifying selection and positive selection (and therefore adaptation) may be more effective in mutualists than non-mutualists because of their larger population sizes.

We took advantage of the presence and absence of nodulation across legumes and rhizobia to test whether mutualistic species evolve more quickly or more slowly than their non-mutualistic relatives. We assessed molecular evolution in 1) six closely related pairs of mutualistic and non-mutualistic plants (i.e., those that do and do not form nodules with rhizobia) across the legume phylogeny, 2) seven pairs of symbiotic and non-symbiotic bacteria species (strains that have *nod* genes and those that lack *nod* genes), and 3) a widespread genus (*Ensifer*) that includes clades of legume symbionts and non-symbiotic bacteria with other lifestyles. We generated *de novo* transcriptomes of 12 non-model legume species to calculate ratios of non-synonymous to synonymous substitutions (dN/dS) at orthologous genes. We also calculated dN/dS values from 14 bacteria species with sequence data deposited in NCBI and from a total of 104 strains in the *Ensifer* phylogeny. We compared dN/dS ratios between mutualistic and non-mutualistic species genome-wide and at symbiotic genes involved in nodulation.

## Methods

### Plant materials and RNA sequencing

We searched several legume phylogenies and clades from the literature (including Zanne et al. 2014 and Azani et al. 2017) and used available nodulation data (Werner et al. 2014) to identify six species pairs of mutualistic legumes and non-mutualistic close relatives. We categorized three of the species pairs as a loss of nodulation because, within these pairs, the non-mutualistic species occurred in a phylogenetic group where at least 90% of the tips were mutualist species. It is unclear whether nodulation has been gained or lost in the other three pairs of species in our analysis. Mutualist species in these pairs are found within a phylogenetic group where 56% of the tips represent non-mutualistic legumes, although this group also includes many tips where the mutualist status is unknown. These pairs could represent nodulation reversals (i.e. a loss followed by a regain), but without more nodulation data, it remains unclear. The six legume species pairs in our analysis are: *Senna italica* and *Senna didymobotrya* (Azani et al. 2017), *Peltophorum africanum* and *Peltophorum dubium* (Haston et al. 2005), *Senna barclayana* and *Senna occidentalis* (LPWG et al. 2013; Azani et al. 2017), *Dalea mollissima* and *Dalea mollis* (McMahon and Hufford 2004; Zanne et al. 2014), *Calliandra eriophylla* and *C. humilis* (Souza et al. 2013), and *Mimosa aculeaticarpa* and *Mimosa grahamii* (Simon et al. 2011) (Fig. 1, Table 1). Due to their shared evolutionary history, comparing closely related species reduces differences between species except for their participation in mutualism. We searched the literature for other traits in the focal species that might influence molecular evolution, including geographic distribution, ploidy, and life history (annual or perennial) (Simonsen et al. 2017; Parshuram et al. 2023), and found that species within pairs generally shared those traits (Table 1). All plants in our dataset likely form indeterminate nodules based on their placement in the legume phylogeny and previous records of indeterminate nodules in the Caesalpinioideae, Mimosoideae, and Papilionoideae legume subfamilies (Andrews and Andrews 2017). We obtained seed for these species from either the USDA-ARS Germplasm Resources Information Network or the KEW Royal Botanical Gardens Millennial Seed Bank.

**Figure 1.**
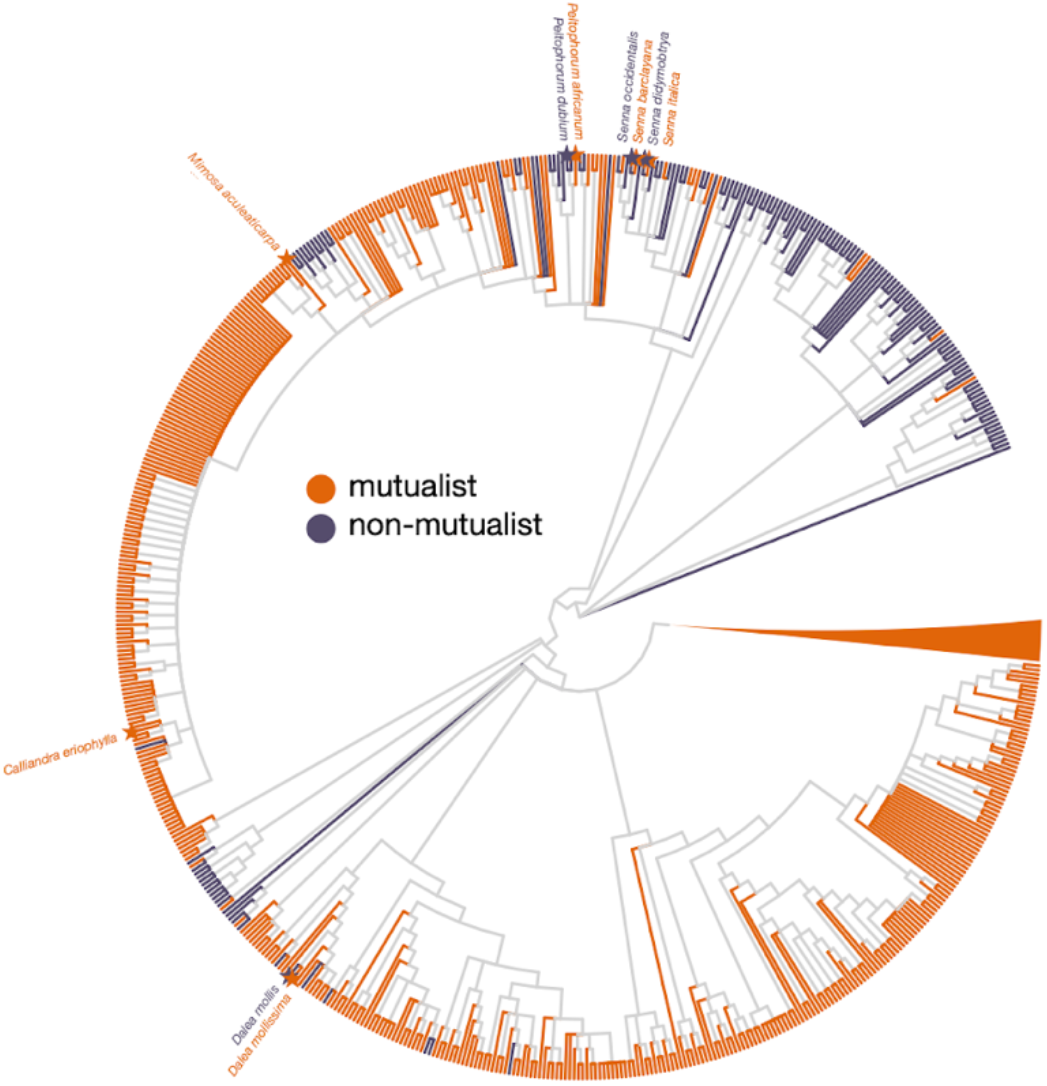
Species pairs of mutualistic (orange) and non-mutualistic legumes (purple) in the study. Sampled legume species are indicated by stars at the tips of the tree and labeled with text. The legume tree developed by the Legume Phylogeny Working Group (LPWG 2013) was filtered for species with symbiotic status data. The branches leading to the tips of the tree were coloured based on whether the species at the tip was known to form nodules (Werner et al. 2014) and therefore is a mutualist (orange) or lacked the ability to nodulate and is therefore a non-mutualist (purple). All internal branches are coloured in grey. The *Calliandra* and *Mimosa* non-symbiotic species were obtained from separate smaller phylogenies, and thus not shown here (Simon et al. 2011; Souza et al. 2013).

**Table 1.** Species of mutualistic and non-mutualistic legumes used in the study. *Senna italica, P. africanum, P. dubium*, and *D. mollis* seeds were sourced from KEW Royal Botanical Gardens Millennial Seed Bank. Seeds of all other species were obtained from USDA-ARS Germplasm Resources Information Network.

| life style | species | introduced ranges | human uses | ploidy | life history | native region |
| --- | --- | --- | --- | --- | --- | --- |
| Non-mutualistic | *Senna didymobotrya* | 20 | 5 | polyploid | perennial | Africa |
| Mutualist | *Senna italica* | 2 | 1 | polyploid | perennial | Africa, Middle East |
| Non-mutualistic | *Peltophorum dubium* | 6 | 3 | polyploid | perennial | South America |
| Mutualist | *Peltophorum africanum* | 7 | 3 | polyploid | perennial | Africa |
| Non-mutualistic | *Senna occidentalis* | 48 | 6 | polyploid | perennial | South America |
| Mutualist | *Senna barclayana* | 0 | 0 | NA | annual/perennial | Australia |
| Non-mutualistic | *Dalea mollis* | 0 | 1 | diploid | annual | North America |
| Mutualist | *Dalea mollissima* | 0 | 1 | diploid | annual | North America |
| Non-mutualistic | *Mimosa grahamii* | 0 | 0 | NA | perennial | North America |
| Mutualist | *Mimosa aculeaticarpa* | 0 | 0 | polyploid | perennial | North/Central America |
| Non-mutualistic | *Calliandra humilis* | 0 | 0 | NA | perennial | North/Central America |
| Mutualist | *Calliandra eriophylla* | 0 | 0 | diploid | perennial | North/Central America |

Because there are no available genomes for our focal legume species, we sequenced RNA from the 12 legume species in our dataset. We grew one plant of each species in a growth chamber with daytime temperature set to 28°C, nighttime temperature set to 19°C, and a light period of 15 hours. We prepared all seeds for germination by nicking the seed coat with a razor blade and incubating the scarified seed at 30°C overnight on wet filter paper in a petri dish. *Senna occidentalis* and *S. barclayana* seeds were placed in boiling water for 10 minutes prior to scarification. *Peltophorum, S. didymobotrya*, and *S. italica* seeds were treated with sulfuric acid for 10 minutes (Alves et al. 2011) and rinsed with distilled water before scarification. All plants were grown in sterile sand in Magenta boxes. Once a week, the bottom compartment of each box was filled with a high-nitrogen fertilizer diluted to one-quarter strength (recipe in Zhang et al. 2020). We did not inoculate plants with rhizobia so that we could collect and compare RNA from roots without nodules from both the mutualistic and non-mutualistic plant species. Previous work has shown that association with rhizobia causes differential expression of many genes with diverse functions, but symbiosis genes are still generally expressed in legumes even in the non-symbiotic state (Afkhami & Stinchcombe 2016). Therefore, symbiosis genes are still captured in transcriptomes from legumes without rhizobia (see Results, below). Plants were harvested for root tissue after five weeks of growth or when the plant had 10 true leaves. Roots were rinsed with water and a small amount of fresh root tissue was cut and stored in Eppendorf tubes. We collected an average of 86 mg of tissue for all species except for the *Peltophorum* species for which we collected 30 mg each. Tubes were flash frozen in liquid nitrogen and stored in a -80°C freezer. We followed the Sigma Aldrich Plant Total RNA Kit instructions to isolate RNA and obtained between 44.1 and 333 ng/ul of RNA per sample. Samples were submitted to Genome Quebec for sequencing on the NovaSeq 6000 Sequencing System (PE100). We received 67,812,540 - 98,924,616 paired end reads per sample with an average quality of 36.

### De novo *transcriptome assembly*

We checked the quality of the sequences with FastQC v0.11.9 (Andrews 2010). We cut adapters, removed leading and trailing low-quality bases (below quality 3), and trimmed sequences to a minimum length of 30 using Trimmomatic v0.39 (Bolger et al. 2014). We assembled *de novo* transcriptomes for each species from the cleaned reads using RNASpades v3.15.2 (Bushmanova et al. 2019) and Trinity v2.8.1 (Haas et al. 2013) with default parameters. We ran CD-HIT v4.8.1 (Fu et al. 2012) to remove redundant transcripts from the assemblies and checked assembly quality with rnaQUEST v2.2.1 (Bushmanova et al. 2019). Transcriptomes produced from RNASpades had fewer but longer contigs, therefore the rest of the analysis was performed on the RNASpades assemblies (Supp. Table 1). We predicted coding regions using TransDecoder v5.5.0 and we removed any contigs with no predicted peptide.

### Rhizobia genome collection

We identified 14 bacteria genomes for analysis by searching rhizobia phylogenies for strains with *nod* genes and close relatives lacking *nod* genes (Rahimlou et al. 2021). We chose seven pairs of nodulating and non-nodulating bacteria species that spanned across both Alpha- and Beta-Proteobacteria. We note that nodulating bacterial species, and their non-nodulating closest relatives, are not the rhizobia partners of the plant species used above. We then downloaded the annotated genomes from NCBI for use in our analysis (Fig. 2a, Table 2). We also downloaded genomes for 65 symbiotic members of the *Ensifer* genus (Fig. 2b, Supp. Table 2) containing *nod* genes and 39 non-symbiotic members without *nod* genes (Fagorzi et al. 2020) for separate analysis comparing symbiotic and non-symbiotic clades.

**Figure 2.**
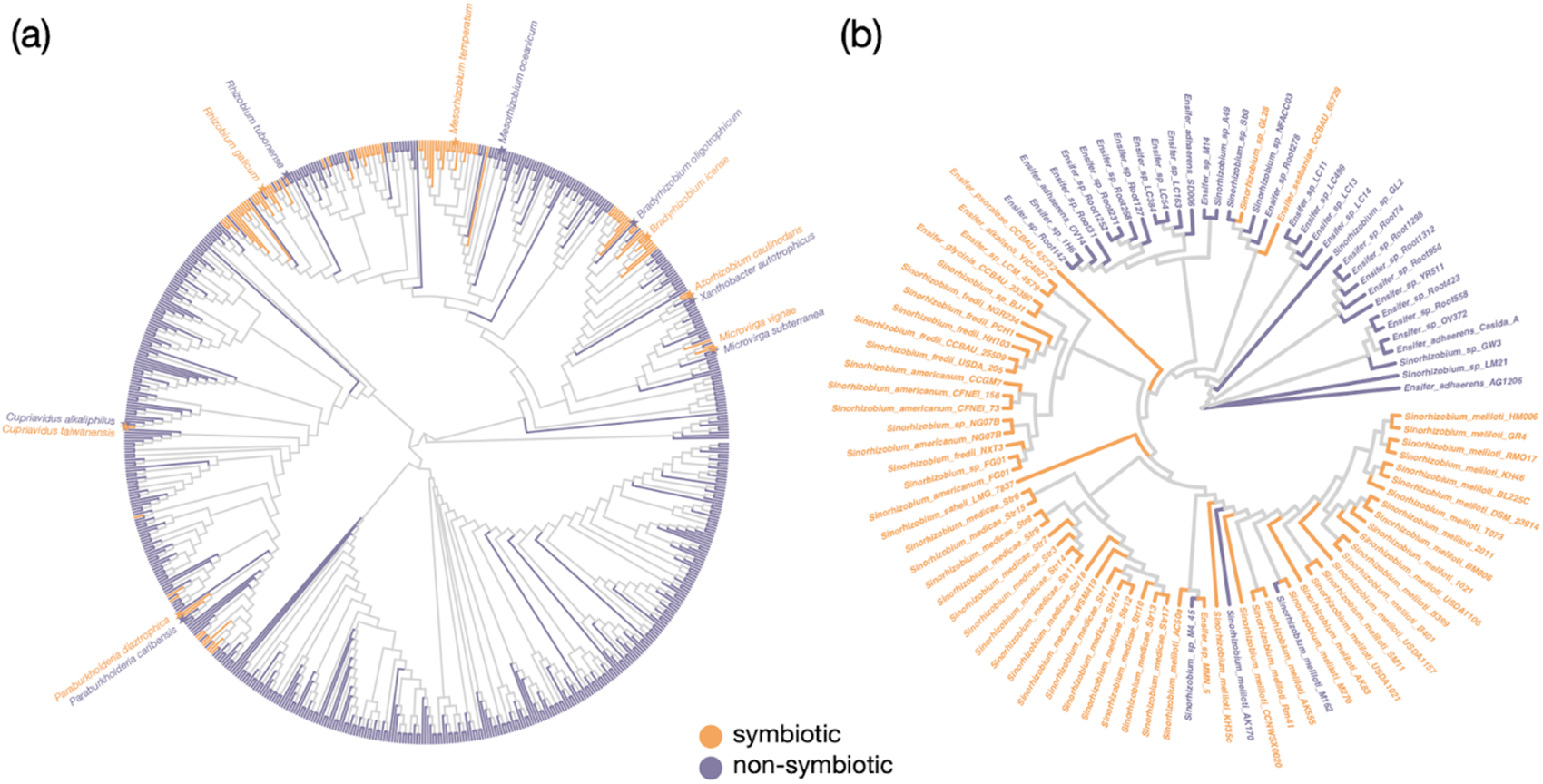
Bacteria species with (orange) or without (purple) *nod* genes. (a) Phylogeny from Rahimlou et al. (2021). The branches leading to the tips of the tree were coloured based on whether the species at the tip is known to have *nod* genes (Rahimlou et al. 2021, Fagorzi et al 2020) and therefore is symbiotic (orange) or lacked *nod* genes and is non-symbiotic (purple). All internal branches are coloured in grey. Species indicated by stars and labeled with text represent the species pairs with genomes used in the analysis. (b) Phylogeny from Fagorzi et al. (2020) trimmed to only the genomes used in the analysis.

**Table 2.** Bacteria strains with genome data collected from NCBI used in the study.

| mutualist | species | assembly code | isolation source |
| --- | --- | --- | --- |
| Non-symbiotic | *Bradyrhizobium oligotrophicum* | GCA_000344805.1 | Paddy field soil |
| Symbiotic | *Bradyrhizobium icense* | GCA_001693385.1 | Root nodule |
| Non-symbiotic | *Xanthobacter autotrophicus* | GCA_000017645.1 | Black sludge |
| Symbiotic | *Azorhizobium caulinodans* | GCA_000010525.1 | Stem nodule |
| Non-symbiotic | *Microvirga subterranea* | GCA_003350535.1 | Geothermal aquifer |
| Symbiotic | *Microvirga vignae* | GCA_001017175.1 | Root nodule |
| Non-symbiotic | *Mesorhizobium oceanicum* | GCA_001889605.1 | Sea water |
| Symbiotic | *Mesorhizobium temperatum* | GCA_002284575.1 | Root nodule |
| Non-symbiotic | *Rhizobium tubonense* | GCA_003240585.1 | Plant |
| Symbiotic | *Rhizobium gallicum* | GCA_001908615.1 | Root nodule |
| Non-symbiotic | *Cupriavidus alkaliphilus* | GCA_003254285.1 | Alkaline soils |
| Symbiotic | *Cupriavidus taiwanensis* | GCA_900250065.1 | Root nodule |
| Non-symbiotic | *Paraburkholderia caribensis* | GCA_001449005.1 | Soil |
| Symbiotic | *Paraburkholderia diazotrophica* | GCA_900108945.1 | Root nodule |

### Ortholog identification

We identified a total of 308 single-copy orthologous genes shared in the proteomes of all 12 legume species using OrthoFinder v2.4 (Emms and Kelly 2019). We expanded this set to also include orthologous genes found in at least four species, resulting in a total of 3548 genes for analysis. We found 438 single-copy orthologous genes in all 14 rhizobia species using default settings in OrthoFinder v2.4 (Emms and Kelly 2019). When we included orthologs present in at least four species, we identified 2812 genes shared among the bacteria strains in our dataset. We identified 456 single copy orthologous genes shared among all 104 *Ensifer* genomes.

### Estimating rates of molecular evolution

To calculate a ratio of nonsynonymous to synonymous substitutions (dN/dS) in each species for each of the 3548 plant orthologous genes and 2812 bacteria genes, we first compiled orthologous nucleotide sequences (cds files) into single fasta files. For each gene, we executed alignments in PRANK v.170427 with the “-codon” option (Markova-Raina and Petrov 2011). We constructed maximum-likelihood gene trees for each orthologous gene using default settings in RAxML with the substitution model set to GTRCATX (Stamatakis 2014). Gene trees and sequence alignments were used as input for dN/dS analysis in PAML v.4.9j (Yang 2007). We implemented a “free-ratios” model in PAML to calculate a separate dN/dS value for each branch in the gene tree. Including six closely related pairs of legumes in the gene trees allows for the pairs to serve as out groups for the different ingroup tests. We extracted dN/dS ratios for each tip of the tree to obtain a unique dN/dS value for each species and performed the remaining analyses in R (R Core Team 2024). We removed all dN/dS ratios greater than 10 as values this high are likely a result of either an error in assembly or overparameterization in the PAML model for complicated gene trees. After filtering abnormally high dN/dS ratios, our sample size included 210 orthologs shared among all 12 plant species and 227 orthologs shared among all 14 bacteria species in our paired analysis. To compare dN/dS ratios between mutualists and non-mutualistic taxa genome-wide, we performed paired Wilcoxon signed-rank tests on dN/dS values for all orthologous genes between mutualist species and their non-mutualistic relative in R (R Core Team 2024; Danneels et al. 2021).

Within species pairs, the plants in our dataset shared similar life history traits. However, across pairs, species differed greatly in their invasion history. We categorized legumes that had been introduced to at least one novel range as invasive and legumes that occur only in their native range as non-invasive. Because invasions may be associated with altered rates of molecular evolution (Whitney & Gabler 2008), we ran a two-rate model in PAML where we allowed all invasive legume species to evolve at one rate and all non-invasive species to evolve at a separate rate. We compared model fit between the two-rate model and a model where all species were constrained to evolve at the same rate. We identified 277 genes out of 308 that showed significant differences in dN/dS ratios between invasive and non-invasive species. We performed paired Wilcoxon signed-rank tests on the significant genes to identify if the invasive genomes are evolving faster than the non-invasive genomes overall.

We also subset our dataset to the three species pairs that represent a loss of mutualism. The species in these pairs are also non-invasive, found in the same habitat (desert habitat in southern USA), and do not play a large role in human agriculture, medicine or industry (genera *Dalea, Mimosa,* and *Calliandra*). For these plant genomes, we implemented a two-ratio model in PAML, where we labeled all species in the gene trees as a mutualist (test) or non-mutualist (reference) and allowed PAML to model separate dN/dS ratios for test and reference branches. We identified the number of genes that showed significantly different dN/dS values in mutualists and non-mutualistic relatives and repeated the paired Wilcoxon signed-rank tests on dN/dS values calculated at these significant genes.

To evaluate rates of molecular in the *Ensifer* genus, we performed a two-ratio model in PAML where we labeled symbiotic species (largely the *Sinorhizobium* clade within the *Ensifer* phylogeny) as the test branches and non-symbiotic species as reference branches. Separate dN/dS ratios were calculated in PAML for test and reference lineages in the *Ensifer* phylogeny. Out of 456 single copy orthologs tested in the two-ratio model, 405 genes showed significant differences between the symbiotic and non-symbiotic clades. We performed paired Wilcoxon signed-rank tests using dN/dS ratios calculated from these 405 significant genes.

### Identifying symbiosis genes

We also compared dN/dS ratios between mutualist and non-mutualist species at genes expected to be involved in the legume-rhizobium symbiosis. As noted above, there are no annotated genomes available for the 12 legume species in our analysis. To identify symbiosis genes in our transcriptomes, we first obtained a list of sequences (Epstein et al. 2022) for 224 genes that Roy et al. (2020) identified as important for symbiosis in *Medicago*. We mapped these sequences to all 12 legume transcriptomes using bwa-mem. We extracted the positions in our transcriptomes where the symbiosis sequences mapped and filtered our full dataset of dN/dS ratios for these genes. We then compared the dN/dS ratios for these symbiosis genes in the mutualist legume and their matching non-symbiotic relative. We also searched for symbiosis genes among the genes that were significantly different in mutualist and non-mutualist species (calculated using two-ratio models in PAML). We performed a blastn search on sequences for any significant symbiosis genes against the flowering plant database (taxid:3398).

To identify symbiosis genes in bacteria, we searched the annotated mutualist genomes for gene descriptions including the following key words: nod, noe, nfe, nodul, nif, fix, fixation, and nitrogenase. Within our set of orthologs, we found 33 genes that contained key words related to nitrogen fixation and nodulation in our dataset of 14 bacteria species. We then compared dN/dS ratios between symbiotic and free-living strains at these symbiosis genes across all pairs in the dataset.

In the *Ensifer* genus, many genes that are important for symbiosis with plants are located within plasmids and symbiotic islands (Geddes et al. 2020). The plasmid pSymB is common among all 104 species in our analysis (Fagorzi et al. 2020), while the presence of pSymA is more variable across strains. Although many *nif* and *nod* genes are located on pSymA (Barnett et al. 2001), there are also a number of symbiosis genes involved in nitrate/nitrite reduction on pSymB (Finan et al. 2001). Therefore, we identified orthologs on pSymB by using the annotated genome assembly for *Sinorhizobium meliloti* USDA1021 (GCA_002197445.1) and analyzed these genes separately from the chromosome and the rest of the genome.

## Results

### Molecular evolution in legumes

We identified 761-1339 matching orthologous genes in the paired legume species. Most pairs showed very similar rates of molecular evolution when we considered genome-wide dN/dS values (Fig. 3). Only one of these comparisons (C. *humilis, C. eriophylla)* showed a significant increase in rates of molecular evolution in the mutualist (Table 3). The other species comparisons showed non-significant differences between mutualists and their non-mutualist relatives.

**Figure 3.**
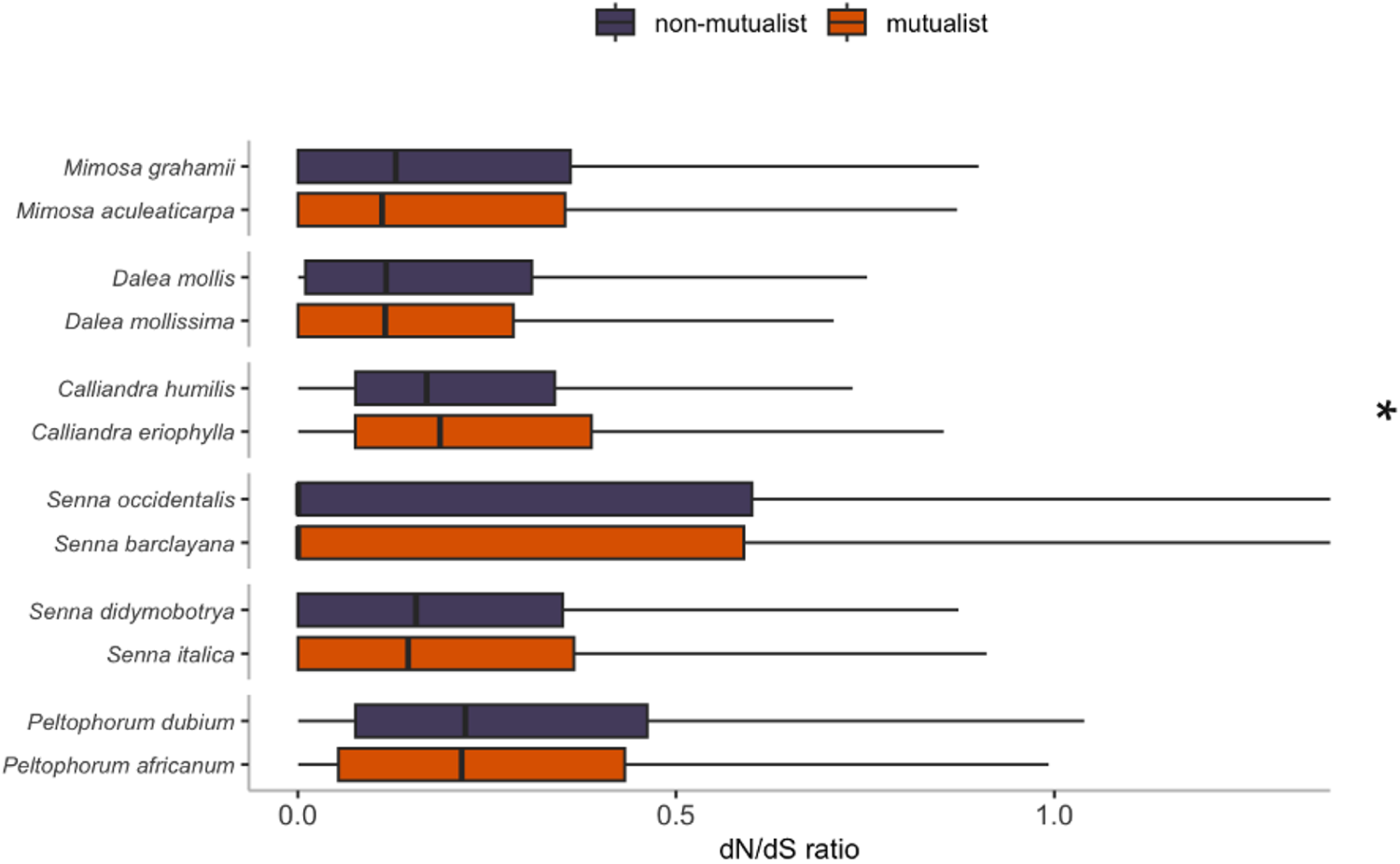
Genome-wide dN/dS ratios estimated from free-ratio models in PAML for mutualistic (orange) and non-mutualistic (purple) legumes. A * indicates species pairs that showed significance at p < 0.05 in paired Wilcoxon tests. Outliers (1.5 x inter quartile range) have been removed from the plot for improved visualization but were included in the paired Wilcoxon signed-rank tests.

**Table 3.** Results of paired Wilcoxon signed-rank tests comparing dN/dS ratios at matching genes in mutualistic legumes and non-mutualistic relatives. The V value is the total sum of ranked genes where the non-mutualistic species had positive values. The U value is the total sum of ranked genes where the mutualist had positive values. The p value is reported for paired Wilcoxon tests where the null hypothesis was that the shift in rank is 0. Significant tests at p<0.05 are bolded.

| mutualist | non-mutualist | gene no. | V | U | p value |
| --- | --- | --- | --- | --- | --- |
| *M. aculeaticarpa* | *M. grahamii* | 1077 | 156945 | 164256 | 0.5769 |
| *D. mollissima* | *D. mollis* | 761 | 109734 | 95386 | 0.1253 |
| ***C. humilis*** | ***C. eriophylla*** | **1191** | **274018** | **329333** | **0.0085** |
| *S. occidentalis* | *S. barclayana* | 1339 | 95093 | 84607 | 0.2160 |
| *S. italica* | *S. didymobotrya* | 1003 | 185751 | 174225 | 0.4193 |
| *P. africanum* | *P. dubium* | 964 | 198736 | 183639 | 0.3120 |

When we analyzed differences between mutualists and non-mutualists in species that are non-invasive and represent a loss of mutualism (using a two-ratio model), we found that mutualist species exhibit increased rates of molecular evolution (Fig. 4a). Directly comparing invasive to non-invasive legumes showed that invasive legumes had higher rates of molecular evolution across the genome (Supp. Fig. 1). The total sum of positive ranked scores for the invasive group was 30761 while the total sum of positive ranked scores for the non-invasive group was 7742 (p<0.001).

**Figure 4.**
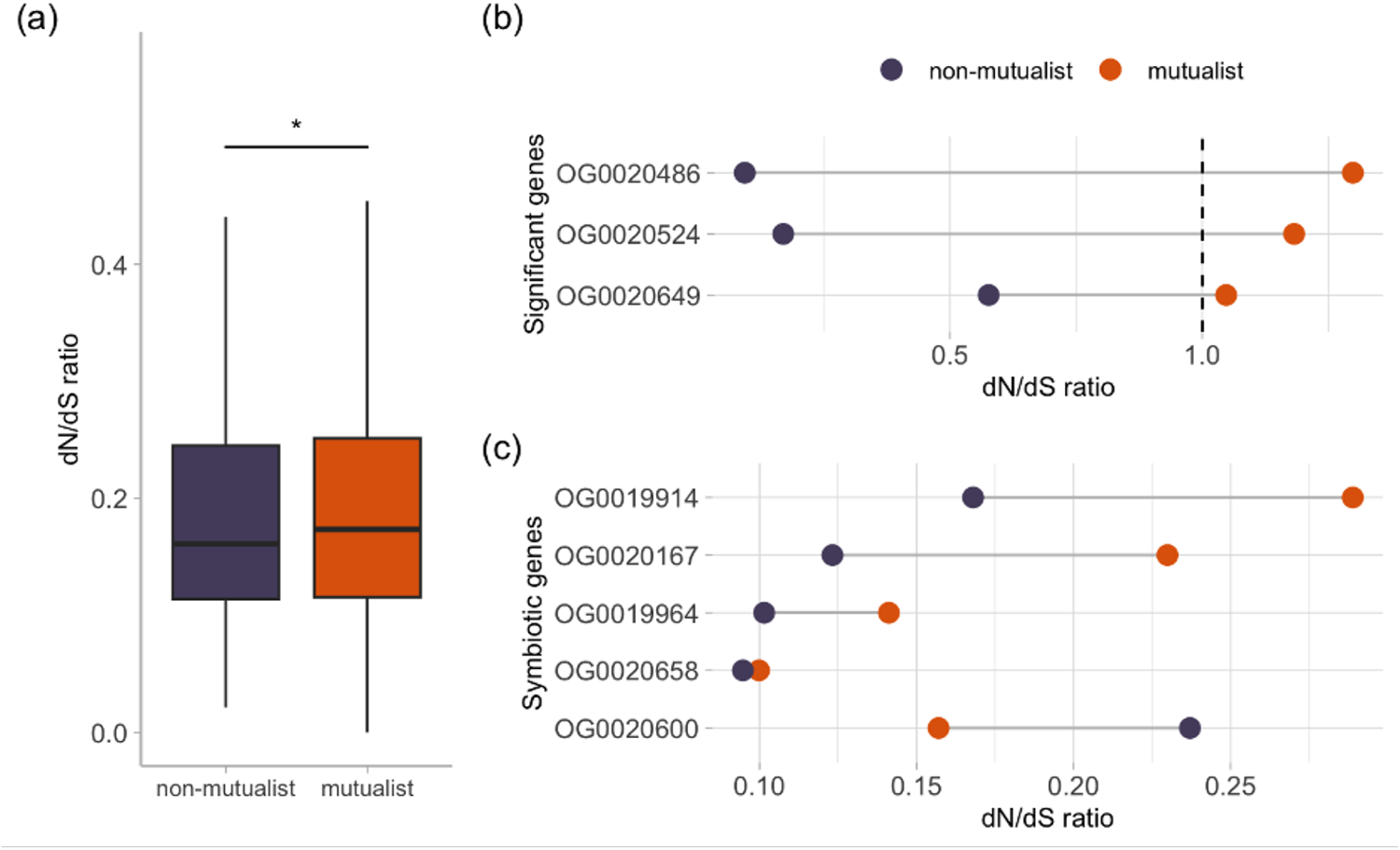
Results from two-ratio models performed in PAML on non-invasive legume species representing a loss of mutualism in the phylogeny. (a) Genome-wide average dN/dS ratios for mutualistic (orange) and non-mutualistic (purple) legumes. A * indicates species pairs that showed significance at p < 0.05 in paired Wilcoxon tests. In this test, a total of 273 genes with significant differences in dN/dS ratios between mutualists and non-symbiotic species (estimated from two-rate PAML models) were included in the analyses. Outliers (1.5 x inter quartile range) have been removed from the plot for improved visualization but were included in the paired Wilcoxon signed-rank tests. (b) Differences in dN/dS ratios for genes under positive selection. (c) Differences in dN/dS ratios for symbiotic genes involved in symbiosis with rhizobia.

### Molecular evolution in rhizobia genomes

We analyzed 836-1288 orthologous genes in the symbiotic and non-symbiotic bacteria pairs. When there were significant differences in evolutionary rates between bacteria species, symbiotic species always had higher dN/dS ratios than non-symbiotic species. Four pairs showed significantly higher rates of molecular evolution in the symbiotic species (Fig. 5). An additional pair, *C. alkaliphilus* and *C. taiwanesis*, also showed higher dN/dS ratios in the symbiotic rhizobia species although this relationship was only marginally significant (Table 4).

**Figure 5.**
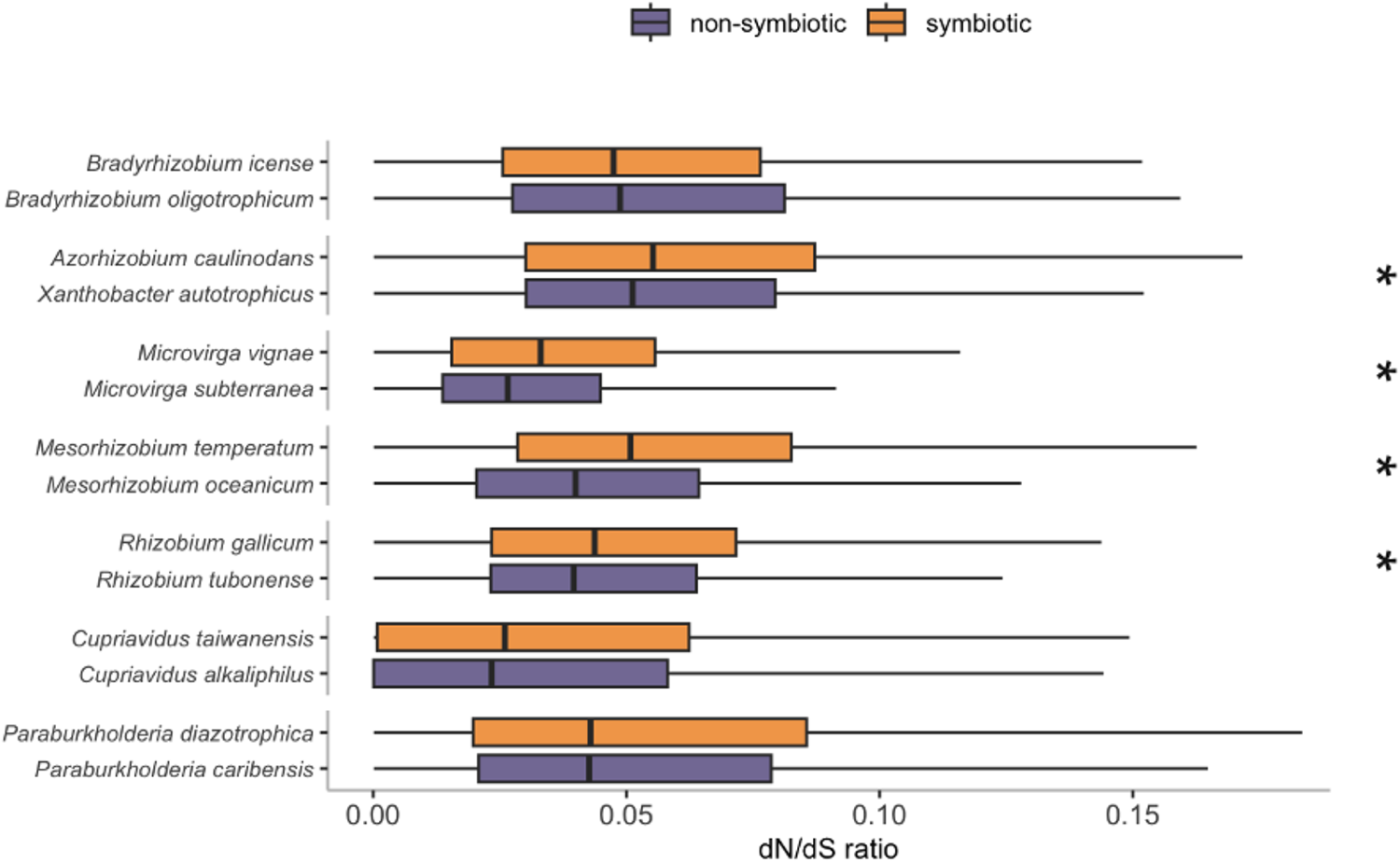
Genome-wide dN/dS ratios estimated from free-ratio models in PAML for symbiotic (orange) and non-symbiotic (purple) bacteria strains. A * indicates species pairs that showed significance at p < 0.05 in paired Wilcoxon tests. Outliers (1.5 x inter quartile range) were removed from the plot for improved visualization but were included in the paired Wilcoxon signed-rank tests.

**Table 4.** Results of paired Wilcoxon signed-rank tests comparing dN/dS ratios at matching genes in symbiotic rhizobia and non-symbiotic relatives. The V value is the total sum of ranked genes where the non-symbiotic species had positive values. The U value is the total sum of ranked genes where the symbiotic species had positive values. The p value is reported for paired Wilcoxon tests where the null hypothesis was that the shift in rank is 0. Significant tests at p<0.05 are bolded.

| **symbiotic** | **non-symbiotic** | **gene no.** | **V** | **U** | **p value** |
| --- | --- | --- | --- | --- | --- |
| *B. icense* | *B. oligotrophicum* | 1154 | 335686 | 326140 | 0.6718 |
| *A. caulinodans* | *X. autotrophicus* | **988** | **200434** | **285171** | **<0.0001** |
| *M. vignae* | *M. subterranea* | **1052** | **216521** | **331060** | **<0.0001** |
| *M. temperatum* | *M. oceanicum* | **1090** | **204387** | **390208** | **<0.0001** |
| *R. gallicum* | *R. tubonense* | **1288** | **373359** | **451611** | **0.0032** |
| *C. taiwanensis* | *C. alkaliphilus* | 836 | 116267 | 136849 | 0.0603 |
| *P. diazotrophica* | *P. caribensis* | 1074 | 262824 | 281622 | 0.3341 |

We analyzed 405 orthologous genes in the comparison between nodulating and non-nodulating *Ensifer* strains. Two-rate models showed that genome wide, the symbiotic strains had higher dN/dS ratios (Fig. 6a, p=0.0003). When we analyzed 70 genes on pSymB separately from the rest of the genome, there was no significant difference between nodulating and non-nodulating strains (Fig. 6b, p=0.2126). Wilcoxon paired tests performed on the chromosome (plus other various accessory plasmids) showed higher dN/dS ratios in the nodulating strains (Fig. 6b, p=0.0007).

**Figure 6.**
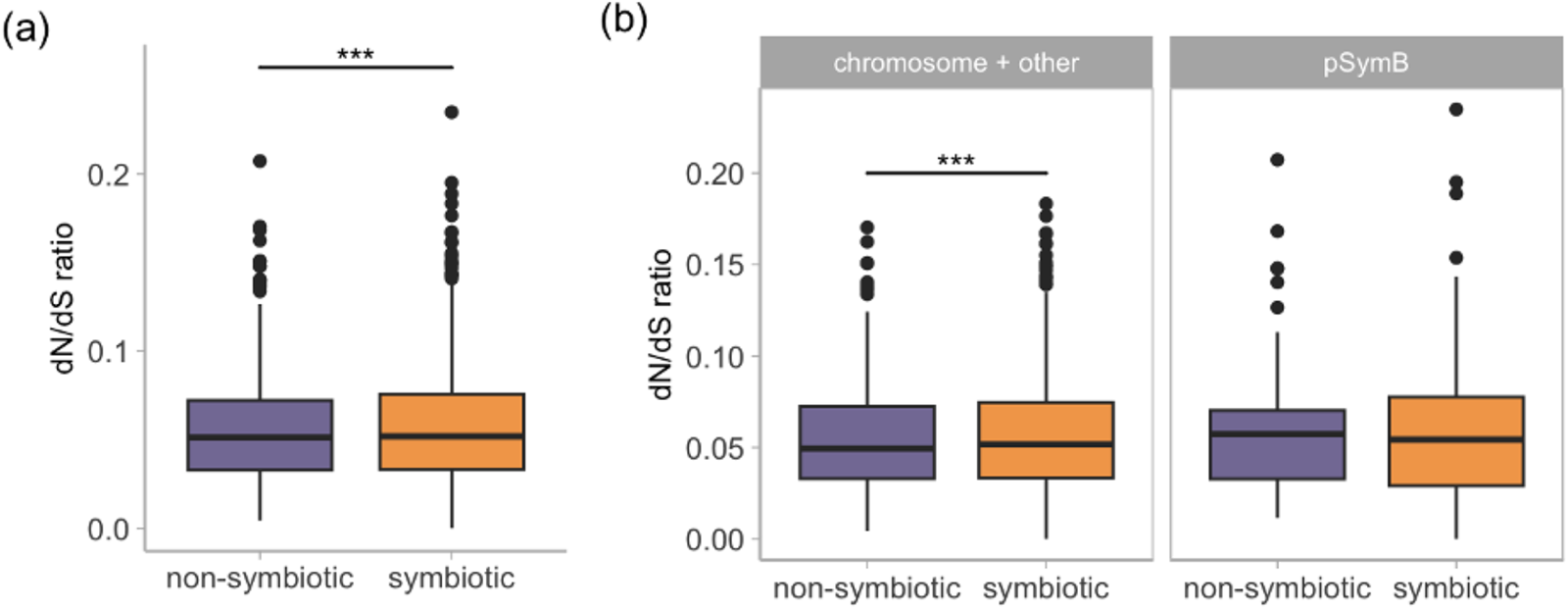
dN/dS ratios estimated from two-rate models in PAML for symbiotic (orange) and non-symbiotic (purple) strains from the *Ensifer* genus. A *** indicates comparisons that showed significance at p < 0.0005 in paired Wilcoxon tests. (a) dN/dS ratios estimated in all genes across the whole genome. (b) dN/dS ratios estimated on genes separated by their location in the genome. The second panel represents dN/dS ratios calculated from genes found on the pSymB plasmid and the first panel shows dN/dS ratios calculated on genes from the rest of the genome including the chromosome and other plasmids (excluding pSymB).

### Symbiosis genes

We identified 17 unique symbiosis genes in our legume ortholog dataset. There was no consistent pattern as to whether these symbiosis genes had higher dN/dS values in the mutualist or non-mutualist plant (Supp. Fig. 2, Supp. Table 3). Differences in dN/dS ratios at symbiosis genes between mutualist and non-mutualistic species were also generally low across the species pairs we tested. One exception is the *Mimosa* pair, where we observed large increases in dN/dS ratios in the non-mutualist *M. grahamii* species compared to the mutualist relative *M. aculeaticarpa* (Supp. Fig. 2).

We identified only five unique symbiosis genes in our filtered dataset for non-invasive legumes when we performed a two-ratio model on these genes in PAML. All but one of these genes had a higher dN/dS value in the mutualist species (Fig. 3c). Only one gene showed an increased rate of molecular evolution in the non-mutualist species. The top hits from the blastn search predicted this gene encodes for a leucine-rich repeat receptor-like serine/threonine-protein kinase.

In the rhizobia genomes, we identified 33 distinct symbiosis genes. For most pairs, there was a fairly equal number of symbiotic and non-symbiotic species with higher dN/dS values at these genes (Supp. Fig. 3, Supp. Table 4). One exception was the non-symbiotic *X. autotrophicus,* which had many more genes with higher rates of molecular evolution (14 genes) compared to its symbiotic relative *A. caulinodans* (5 genes).

### Genes under positive selection

Using the free-ratio dataset, we identified a total of 797 unique genes that were under positive selection in plant species. The number of genes under positive selection varied across species pairs (Supp. Table 5). The dN/dS ratios at these genes were not consistently higher in one species over the other. When we subset our dataset to genes found in all 12 species, we found 78 genes under positive selection. Few of these genes were under positive selection in more than one species in our dataset (Supp. Fig. 4). We found only three genes under positive selection when we considered non-invasive legume dN/dS ratios calculated from two-ratio models in PAML (Fig. 4b). All three genes had higher dN/dS ratios in the mutualist species and a dN/dS ratio under 1 in the non-mutualistic relatives. We identified these genes as a telomere repeat-binding factor, very-long-chain (3R)-3-hydroxyacyl-CoA dehydratase, and 50S ribosomal chloroplastic protein using blastn searches.

In rhizobia, we found 110 unique genes across all pairs that were under positive selection. There was no consistent pattern as to whether symbiotic species or non-symbiotic species contained more genes under positive selection (Supp. Table 6). When we considered genes that were common to all 14 rhizobia genomes, only 13 were under positive selection. None of these genes had dN/dS ratios greater than one in multiple species (Supp. Fig. 5).

## Discussion

In this study, we investigated shifts in rates of molecular evolution genome-wide and at symbiotic genes in 1) mutualistic versus non-mutualistic legumes, 2) symbiotic versus non-symbiotic rhizobia, and 3) symbiotic and non-symbiotic clades in the *Ensifer* phylogeny. We sequenced and assembled transcriptomes for 12 non-model plant species from across the legume phylogeny. In bacteria, we collected rhizobia genomes from across the Alpha- and Beta-Proteobacteria clades and analyzed 104 genomes across the *Ensifer* phylogeny. When there were significant differences in rates of molecular evolution between mutualists and non-mutualistic species, mutualists always showed faster evolutionary rates genome-wide. When we examined symbiosis genes in both legumes and rhizobia, mutualists did not consistently show higher dN/dS values compared to non-mutualist species. We consider in turn several possible explanations for faster evolutionary rates genome-wide, but not at symbiosis genes, in mutualist legumes and rhizobia.

The first potential explanation we consider is coevolution between legumes and rhizobia. If legumes and rhizobia are engaged in ongoing coevolution (i.e. reciprocal adaptation and response), we might predict that increased positive selection would elevate rates of molecular evolution, as reported in parasitic systems (Paterson et al. 2010; Bromham et al. 2013). However, if species are coevolving, we would also expect to see especially high dN/dS ratios at symbiosis genes in mutualist lineages. All the symbiosis genes in our dataset were under purifying selection in both legumes and rhizobia (except for one gene in the *Mimosa* genus), and it was variable whether rates were relatively higher or lower in the mutualist partner. Only our analyses on legumes that have apparently lost nodulation showed that mutualists consistently had higher dN/dS ratios than non-mutualistic legumes at symbiotic genes (though still less than one). Previous work has also failed to find strong evidence for population genetic signatures of balancing or positive selection driving evolution of symbiosis genes in legumes (Yoder 2016; Epstein et al. 2022), consistent with our results.

It is possible that different legumes may not have evolved the same symbiosis genes to associate with rhizobia and by mapping known symbiosis genes in *Medicago* to our transcriptomes, we may be missing some key symbiosis genes in our non-model legume species. We found little overlap in genes under positive selection across the dataset, also suggesting that different species are experiencing different selective pressures targeting different genes. However, some symbiosis genes in legumes are highly conserved (Schnabel et al. 2011; Roy et al. 2020) and therefore unlikely to show signs of positive selection expected from coevolution. Symbiosis genes in rhizobia also show evolutionary conservation (Laranjo et al. 2008) and the ability to nodulate plants is largely acquired through the horizontal transfer of symbiotic plasmids (Wernegreen and Riley 1999) or symbiotic islands (Sullivan et al. 1995), providing further support for this interpretation. One hypothetical possibility is that after accepting a symbiotic plasmid or island, a bacterium may undergo many mutations in the rest of the genome to accommodate the new genetic material that allows it access to a plant host. Previous research suggests that the initial introduction of plasmids with symbiotic genes is not enough to maintain cooperation in bacteria long term (Dewar et al. 2021). Therefore, a large-scale change to the genome plus living in a new habitat (root nodule) may drive up dN/dS ratios across the chromosome, but not at genes on the symbiotic plasmid itself.

There could be other mechanisms driving differences in substitution rates between species other than mutualism. Differences in life history strategies in plants have been previously shown to influence rates of molecular evolution. For instance, the generation time hypothesis predicts that long-lived species (i.e. perennials) evolve more slowly compared to annuals (Smith and Donoghue 2008). Additionally, asexual plant species show increased accumulation of substitutions compared to sexually reproducing species (Hollister et al. 2015). Duplicated genes are likely to experience higher rates of molecular evolution (Kimura and Ohta 1974), indicating that organisms with higher ploidy levels might show elevated dN/dS values. Mutualists and non-mutualists generally had similar life history traits (ploidy level, generation time, reproduction) within species pairs in our datasets. Therefore, whether plants participate in mutualism with rhizobia should be the main lifestyle difference between species in our dataset. Our analyses also showed that invasive legumes evolve at a faster rate compared to non-invasive species. High rates of molecular evolution may make plants better at invading new habitats as fast evolving organisms could have greater niche breadth and environmental tolerances. Alternatively, elevated rates of molecular evolution may be a consequence of invasion because once established in a new environment, plants may have to adapt quickly (Young et al. 2018). However, short evolutionary timeframes post invasion may not be long enough for plant genomes to accumulate substitutions. While we cannot account for all possible differences in life history strategies (or other traits) between species, we were able to match most life history traits within species pairs when we could find data on these key traits in the literature. Therefore, differences in molecular evolution between mutualists and non-mutualists in our analyses are unlikely to be fully explained by differences in life history traits.

Another potential class of explanations for our results is the efficacy of selection in mutualist populations. Population size is predicted to have drastic effects on genome and molecular evolution. Species with small population sizes are expected to accumulate more deleterious mutations due to genetic drift (Charlesworth 2009). Previous work comparing island to mainland species (Woolfit and Bromham 2005) and small mammals to large mammals (Popadin et al. 2007) has shown that species with small population sizes experience faster rates of molecular evolution. In contrast, genetic drift is less pronounced in large populations and selection is more efficient (Charlesworth 2009). Species engaged in mutualism are expected to grow to larger population sizes because having a beneficial partner can help organisms occupy novel habitats, access nutrients when resources are scarce, and resist natural enemies (Afkhami et al. 2014; Weber and Agrawal 2014; Hayward et al. 2015). Therefore, the high dN/dS ratios in mutualists might be a result of large mutualist populations experiencing (more) positive selection than non-mutualists. Alternatively, higher dN/dS ratios across mutualist genomes could be a result of relaxed negative selection (Bromham et al. 2013). If mutualist legumes always rely on a rhizobia partner for access to nitrogen, there may be less selective pressure to maintain other genes that are important for accessing nutrients in the absence of a rhizobia partner. For instance, genes responsible for root proliferation might be under relaxed selection if it is less important for plants to ‘forage’ for soil nutrients when rhizobia are present. Previous work has shown that mutualist traits and root foraging show a weak (quantitative) genetic correlation (Batstone et al. 2017), suggesting that traits could be evolving largely independently (*i.e*., relaxed selection on root foraging genes while purifying selection acts on symbiotic genes). Given that symbiotic genes in rhizobia are clustered on plasmids or in genomic islands (Wernegreen and Riley 1999), there is also opportunity for the chromosome to experience relaxed selection while purifying selection simultaneously acts on symbiotic plasmids or islands in bacteria. In addition, if rhizobia undergo more replication events while inside nodules than in the soil, then there could be relaxed selection on genes encoded on the chromosome for traits that only matter in the soil environment (*e.g.*, competition with other microbes) and not inside the intracellular nodule environment.

## Conclusion

In our study, genome-wide elevated rates of molecular evolution is a common feature of both mutualist partners in the legume-rhizobium symbiosis. Genetic analyses of other positive species interactions also show accelerated rates of molecular evolution in mutualists species, suggesting that our findings are a general characteristic of mutualism overall (Lutzoni and Pagel 1997; Rubin and Moreau 2016). Slower evolution was particularly evident in three non-mutualistic legumes that appear to have lost nodulation. In both plants and rhizobia, there was no consistent pattern in rates of molecular evolution at symbiotic genes which were largely under purifying selection. A combination of relaxed selection and more effective positive selection in large mutualist populations may be responsible for the high rates of molecular evolution we observed in mutualist legumes and rhizobia.

## Data availability

Sequence data will be uploaded to SRA upon acceptance. Assembly codes for the bacteria and rhizobia genomes used in the analysis are listed in Table 2 and Supplemental Table 2. All code for reproducing the analysis will be made public on a github repository following acceptance for publication.

## Acknowledgements

We thank Stephen Wright, Mark Hibbins, and Tyler Kent for advice on running PAML models and Matt Pennell for suggestions on analyzing dN/dS ratios. We also thank Maria Tocora for advice on running transcriptome assembly programs and George diCenzo for feedback on analyzing the *Ensifer* samples. Our work is supported by NSERC and Queen Elizabeth II/Charles E. Eckenwalder Graduate Scholarships (TLH) and NSERC Discovery Grants (JRS, MEF).

