## Supplemental Materials for "Elevated rates of molecular evolution genome-wide in mutualist legumes and rhizobia"

### Supplemental Information

**Supplemental Table 1. Summary statistics for *de novo* transcriptome assemblies.** Reported contig numbers are after collapsing transcripts. ORFs are open reading frames predicted by TransDecoder.

| **RNASpades assemblies** | | | | | |
|---|---|---|---|---|---|
| **Sample** | **Transcript no.** | **Contig no.** | **Ave. transcript length** | **N50** | **ORFs no.** |
| SD2 _1 | 68820807 | 117982 | 1123.91 | 2108 | 61303 |
| SI2_1 | 98924616 | 112317 | 1306.248 | 2375 | 66365 |
| DA2 | 80577865 | 150995 | 722.742 | 1321 | 56375 |
| DS2 | 75572333 | 107221 | 1177.985 | 2240 | 56669 |
| PA1_2 | 76087393 | 101736 | 1320.368 | 2284 | 59500 |
| PD1_4 | 71804897 | 102658 | 1273.57 | 2282 | 58082 |
| SO2 | 89355436 | 130849 | 1154.002 | 2287 | 63787 |
| SB2 | 67812540 | 126102 | 1009.44 | 2015 | 56847 |
| MA4 | 64080579 | 160887 | 1035.987 | 1857 | 80753 |
| MG1 | 76797627 | 136567 | 1098.058 | 2027 | 68972 |
| CH1 | 81835443 | 100378 | 1313.984 | 2304 | 62339 |
| CE2 | 77242582 | 158564 | 1017.964 | 1861 | 82106 |
| **Trinity assemblies** | | | | | |
| SD2_1 | 68820807 | 171246 | 1090.601 | 2067 | |

| | | | | |
|---|---|---|---|---|
| SI2_1 | 98924616 | 171274 | 1233.415 | 2259 |
| DA2 | 80577865 | 168634 | 732.721 | 1338 |
| DS2 | 75572333 | 149461 | 1145.836 | 2163 |
| PA1_2 | 76087393 | 158493 | 1289.341 | 2231 |
| PD1_4 | 71804897 | 153582 | 1289.253 | 2285 |
| SO2 | 89355436 | 195385 | 1111.781 | 2187 |
| SB2 | 67812540 | 176262 | 1032.021 | 2030 |
| MA4 | 64080579 | 252925 | 917.198 | 1643 |
| MG1 | 76797627 | 205855 | 995.444 | 1866 |
| CH1 | 81835443 | 158130 | 1220.887 | 2204 |
| CE2 | 77242582 | 233601 | 899.113 | 1691 |

**Supplemental Table 2. Genome assemblies used in analyses on the *Ensifer* genus.**

| organism | assembly code | assembly level | mutualist |
|---|---|---|---|
| *Ensifer adhaerens* | GCA_000697965.2 | Complete | Non-symbiotic |
| *Ensifer adhaerens* | GCA_001270265.1 | Contig | Non-symbiotic |
| *Ensifer adhaerens* | GCA_003269115.1 | Scaffold | Non-symbiotic |
| *Ensifer adhaerens* | GCA_013283195.1 | Scaffold | Non-symbiotic |
| *Ensifer adhaerens* | GCA_000583045.1 | Chromosome | Non-symbiotic |
| *Ensifer alkalisoli* | GCA_001723275.1 | Scaffold | Symbiotic |
| *Ensifer glycinis* | GCA_001651865.1 | Contig | Symbiotic |
| *Ensifer psoraleae* | GCA_013283645.1 | Scaffold | Symbiotic |
| *Ensifer sesbaniae* | GCA_013283665.1 | Scaffold | Symbiotic |
| *Ensifer sp* | GCA_001976035.1 | Contig | Non-symbiotic |
| *Ensifer sp* | GCA_001695835.1 | Contig | Non-symbiotic |
| *Ensifer sp* | GCA_001695795.1 | Contig | Non-symbiotic |
| *Ensifer sp* | GCA_001695785.1 | Contig | Non-symbiotic |
| *Ensifer sp* | GCA_001695855.1 | Contig | Non-symbiotic |
| *Ensifer sp* | GCA_001695865.1 | Contig | Non-symbiotic |
| *Ensifer sp* | GCA_001695895.1 | Contig | Non-symbiotic |
| *Ensifer sp* | GCA_001695905.1 | Contig | Non-symbiotic |
| *Ensifer sp* | GCA_001854885.1 | Contig | Symbiotic |
| *Ensifer sp* | GCA_003355565.1 | Scaffold | Non-symbiotic |
| *Ensifer sp* | GCA_002885935.1 | Contig | Symbiotic |
| *Ensifer sp* | GCA_003024455.1 | Contig | Non-symbiotic |
| *Ensifer sp* | GCA_900113205.1 | Scaffold | Non-symbiotic |
| *Ensifer sp* | GCA_001425885.1 | Scaffold | Non-symbiotic |
| *Ensifer sp* | GCA_001425225.1 | Scaffold | Non-symbiotic |
| *Ensifer sp* | GCA_001426365.1 | Contig | Non-symbiotic |
| *Ensifer sp* | GCA_001426465.1 | Contig | Non-symbiotic |

| | | | |
|---|---|---|---|
| *Ensifer sp* | GCA_001426785.1 | Scaffold | Non-symbiotic |
| *Ensifer sp* | GCA_001428785.1 | Scaffold | Non-symbiotic |
| *Ensifer sp* | GCA_001429745.1 | Scaffold | Non-symbiotic |
| *Ensifer sp* | GCA_001429285.1 | Scaffold | Non-symbiotic |
| *Ensifer sp* | GCA_001424825.1 | Scaffold | Non-symbiotic |
| *Ensifer sp* | GCA_001426275.1 | Scaffold | Non-symbiotic |
| *Ensifer sp* | GCA_001427045.1 | Scaffold | Non-symbiotic |
| *Ensifer sp* | GCA_001429125.1 | Contig | Non-symbiotic |
| *Ensifer sp* | GCA_001429005.1 | Scaffold | Non-symbiotic |
| *Ensifer sp* | GCA_900103045.1 | Scaffold | Non-symbiotic |
| *Sinorhizobium americanum* | GCA_001651855.1 | Contig | Symbiotic |
| *Sinorhizobium americanum* | GCA_001889105.1 | Complete | Symbiotic |
| *Sinorhizobium americanum* | GCA_002909045.1 | Contig | Symbiotic |
| *Sinorhizobium americanum* | GCA_002909075.1 | Contig | Symbiotic |
| *Sinorhizobium americanum* | GCA_000705595.2 | Complete | Symbiotic |
| *Sinorhizobium fredii* | GCA_002531965.1 | Contig | Symbiotic |
| *Sinorhizobium fredii* | GCA_002944405.1 | Complete | Symbiotic |
| *Sinorhizobium fredii* | GCA_003177055.1 | Complete | Symbiotic |
| *Sinorhizobium fredii* | GCA_000283895.1 | Chromosome | Symbiotic |
| *Sinorhizobium fredii* | GCA_000018545.1 | Complete | Symbiotic |
| *Sinorhizobium fredii* | GCA_001461695.1 | Contig | Symbiotic |
| *Sinorhizobium medicae* | GCA_002864945.1 | Scaffold | Symbiotic |
| *Sinorhizobium medicae* | GCA_002864955.1 | Scaffold | Symbiotic |
| *Sinorhizobium medicae* | GCA_002864985.1 | Scaffold | Symbiotic |
| *Sinorhizobium medicae* | GCA_002865005.1 | Scaffold | Symbiotic |
| *Sinorhizobium medicae* | GCA_002865025.1 | Scaffold | Symbiotic |
| *Sinorhizobium medicae* | GCA_002865035.1 | Scaffold | Symbiotic |
| *Sinorhizobium medicae* | GCA_002865045.1 | Scaffold | Symbiotic |

| | | | |
|---|---|---|---|
| *Sinorhizobium medicae* | GCA_002865055.1 | Scaffold | Symbiotic |
| *Sinorhizobium medicae* | GCA_002865105.1 | Scaffold | Symbiotic |
| *Sinorhizobium medicae* | GCA_002865125.1 | Scaffold | Symbiotic |
| *Sinorhizobium medicae* | GCA_002865145.1 | Scaffold | Symbiotic |
| *Sinorhizobium medicae* | GCA_002865155.1 | Scaffold | Symbiotic |
| *Sinorhizobium medicae* | GCA_002865185.1 | Scaffold | Symbiotic |
| *Sinorhizobium medicae* | GCA_002865225.1 | Scaffold | Symbiotic |
| *Sinorhizobium medicae* | GCA_002865265.1 | Scaffold | Symbiotic |
| *Sinorhizobium medicae* | GCA_000017145.1 | Complete | Symbiotic |
| *Sinorhizobium meliloti* | GCA_000747295.1 | Complete | Symbiotic |
| *Sinorhizobium meliloti* | GCA_000968555.1 | Contig | Symbiotic |
| *Sinorhizobium meliloti* | GCA_002197025.1 | Complete | Symbiotic |
| *Sinorhizobium meliloti* | GCA_002197065.1 | Complete | Symbiotic |
| *Sinorhizobium meliloti* | GCA_002197085.1 | Complete | Symbiotic |
| *Sinorhizobium meliloti* | GCA_002197105.1 | Complete | Symbiotic |
| *Sinorhizobium meliloti* | GCA_002197125.1 | Complete | Non-symbiotic |
| *Sinorhizobium meliloti* | GCA_002197145.1 | Complete | Symbiotic |
| *Sinorhizobium meliloti* | GCA_002197165.1 | Complete | Symbiotic |
| *Sinorhizobium meliloti* | GCA_002197445.1 | Complete | Symbiotic |
| *Sinorhizobium meliloti* | GCA_002197465.1 | Complete | Symbiotic |
| *Sinorhizobium meliloti* | GCA_002215195.1 | Complete | Symbiotic |
| *Sinorhizobium meliloti* | GCA_002302355.1 | Complete | Symbiotic |
| *Sinorhizobium meliloti* | GCA_002302375.1 | Complete | Symbiotic |
| *Sinorhizobium meliloti* | GCA_002807095.1 | Scaffold | Symbiotic |
| *Sinorhizobium meliloti* | GCA_003034185.1 | Scaffold | Symbiotic |
| *Sinorhizobium meliloti* | GCA_003044175.2 | Contig | Symbiotic |
| *Sinorhizobium meliloti* | GCA_003044215.2 | Contig | Symbiotic |
| *Sinorhizobium meliloti* | GCA_003692735.1 | Scaffold | Non-symbiotic |

| | | | |
|---|---|---|---|
| *Sinorhizobium meliloti* | GCA_900107055.1 | Scaffold | Symbiotic |
| *Sinorhizobium meliloti* | GCA_900108935.1 | Scaffold | Symbiotic |
| *Sinorhizobium meliloti* | GCA_000006965.1 | Complete | Symbiotic |
| *Sinorhizobium meliloti* | GCA_000346065.1 | Complete | Symbiotic |
| *Sinorhizobium meliloti* | GCA_000147775.3 | Complete | Symbiotic |
| *Sinorhizobium meliloti* | GCA_000236945.2 | Contig | Symbiotic |
| *Sinorhizobium meliloti* | GCA_000320385.2 | Complete | Symbiotic |
| *Sinorhizobium meliloti* | GCA_000304415.1 | Complete | Symbiotic |
| *Sinorhizobium meliloti* | GCA_001050915.2 | Complete | Symbiotic |
| *Sinorhizobium meliloti* | GCA_000218265.1 | Complete | Symbiotic |
| *Sinorhizobium saheli* | GCA_001651875.1 | Scaffold | Symbiotic |
| *Sinorhizobium sp* | GCA_002002725.1 | Contig | Non-symbiotic |
| *Sinorhizobium sp* | GCA_002531555.1 | Contig | Symbiotic |
| *Sinorhizobium sp* | GCA_002531875.1 | Contig | Symbiotic |
| *Sinorhizobium sp* | GCA_001461715.1 | Contig | Non-symbiotic |
| *Sinorhizobium sp* | GCA_001461685.1 | Contig | Symbiotic |
| *Sinorhizobium sp* | GCA_001461765.1 | Contig | Non-symbiotic |
| *Sinorhizobium sp* | GCA_002216665.1 | Contig | Non-symbiotic |
| *Sinorhizobium sp* | GCA_002885915.1 | Contig | Non-symbiotic |
| *Sinorhizobium sp* | GCA_900103435.1 | Scaffold | Non-symbiotic |
| *Sinorhizobium sp* | GCA_002532005.1 | Contig | Symbiotic |
| *Sinorhizobium sp* | GCA_001461705.1 | Contig | Non-symbiotic |

**Supplemental Table 3. Summary statistics of dN/dS ratios on legume symbiotic genes calculated from free-ratio models in PAML.** The number of genes where the mutualist had a higher value (M higher), where the non-mutualistic species had a higher value (N higher) and when the gene had the same value in both species are reported.

| mutualist (m) | non-mutualist (n) | gene no | m higher | n higher | equal |
|---|---|---|---|---|---|
| *M. aculeaticarpa* | *M grahamii* | 11 | 3 | 7 | 1 |
| *D. mollissima* | *D. mollis* | 9 | 6 | 3 | 0 |
| *C. humilis* | *C. eriophylla* | 12 | 8 | 4 | 0 |
| *S. occidentalis* | *S. barclayana* | 10 | 1 | 2 | 7 |
| *S. italica* | *S. didymobotrya* | 8 | 4 | 4 | 0 |
| *P. africanum* | *P. dubium* | 11 | 6 | 4 | 0 |

**Supplemental Table 4. Summary statistics of dN/dS ratios on rhizobia symbiotic genes calculated from free-ratio models in PAML.** The number of genes where the symbiotic species had a higher value (M higher), where the non-symbiotic species had a higher value (F higher) and when the gene had the same value in both species are reported.

| **symbiotic (s)** | **non-symbiotic (s)** | **gene no** | **s higher** | **n higher** | **equal** |
|---|---|---|---|---|---|
| *B. icense* | *B. oligotrophicum* | 22 | 12 | 10 | 0 |
| *A. caulinodans* | *X. autotrophicus* | 19 | 5 | 14 | 0 |
| *M. vignae* | *M. subterranea* | 7 | 3 | 4 | 0 |
| *M. temperatum* | *M. oceanicum* | 9 | 5 | 4 | 0 |
| *R. gallicum* | *R. tubonense* | 8 | 1 | 7 | 0 |
| *C. taiwanensis* | *C. alkaliphilus* | 7 | 5 | 2 | 0 |
| *P. diazotrophica* | *P. caribensis* | 9 | 5 | 4 | 0 |

**Supplemental Table 5. Summary statistics of dN/dS ratios on legume genes under positive selection calculated from free-ratio models in PAML.** The number of genes where the mutualist had a higher value (M higher), where the non-mutualist species had a higher value (N higher) and when the gene had the same value in both species are reported. A total of 797 unique genes were found to be under positive selection across all species.

| mutualist (m) | non-mutualistic (n) | gene no | m higher | n higher |
|---|---|---|---|---|
| *M. aculeaticarpa* | *M grahamii* | 164 | 90 | 74 |
| *D. mollissima* | *D. mollis* | 89 | 31 | 58 |
| *C. humilis* | *C. eriophylla* | 142 | 92 | 60 |
| *S. occidentalis* | *S. barclayana* | 293 | 141 | 152 |
| *S. italica* | *S. didymobotrya* | 98 | 40 | 58 |
| *P. africanum* | *P. dubium* | 128 | 67 | 61 |

**Supplemental Table 6. Summary statistics of dN/dS ratios on rhizobia genes under positive selection calculated from free-ratio models in PAML.** The number of genes where the symbiotic species had a higher value (M higher), where the non-symbiotic species had a higher value (F higher) and when the gene had the same value in both species are reported.

| symbiotic (s) | non-symbiotic (n) | gene no | s higher | n higher |
|---|---|---|---|---|
| *B. icense* | *B. oligotrophicum* | 3 | 2 | 1 |
| *A. caulinodans* | *X. autotrophicus* | 6 | 3 | 3 |
| *M. vignae* | *M. subterranea* | 3 | 3 | 0 |
| *M. temperatum* | *M. oceanicum* | 4 | 2 | 2 |
| *R. gallicum* | *R. tubonense* | 3 | 2 | 1 |
| *C. taiwanensis* | *C. alkaliphilus* | 45 | 23 | 22 |
| *P. diazotrophica* | *P. caribensis* | 25 | 13 | 12 |

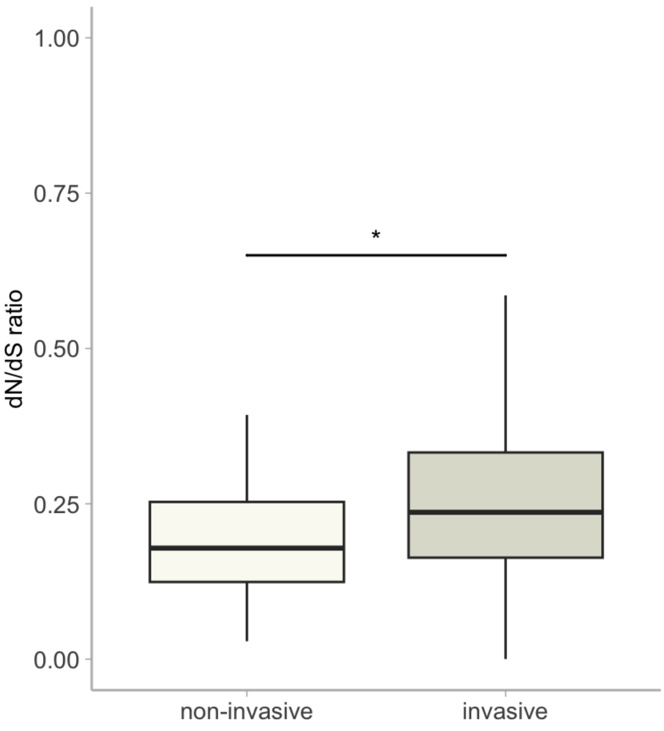


**Supplemental Figure 1. Relationship between dN/dS values and invasive status of legumes.** Significance based on linear model results are indicated by * $p<0.05$.

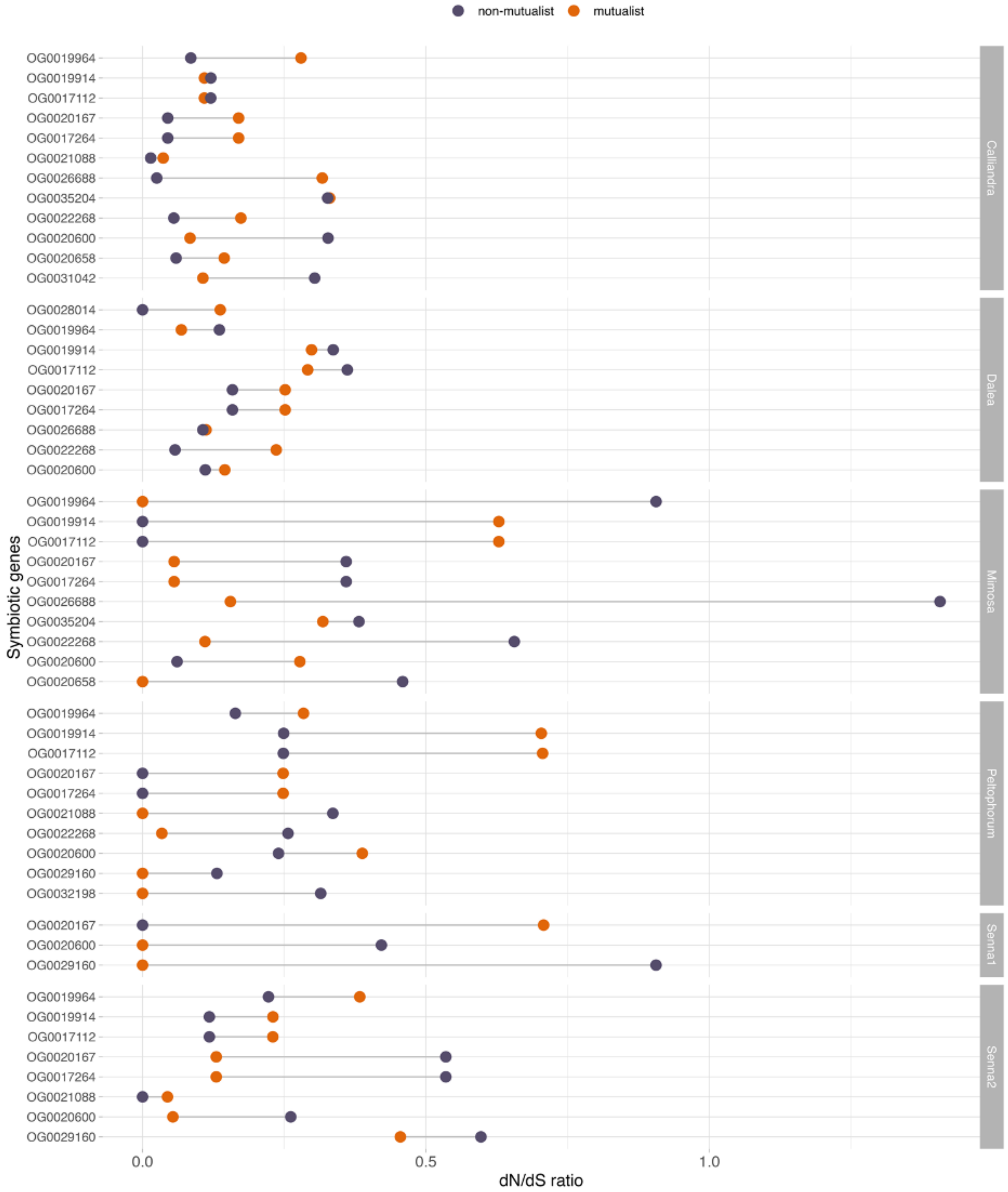


**Supplemental Figure 2. Differences in dN/dS ratios between mutualistic (orange) and non-mutualistic (purple) plant species at symbiotic genes.** dN/dS ratios were calculated from free-ratio models in PAML.

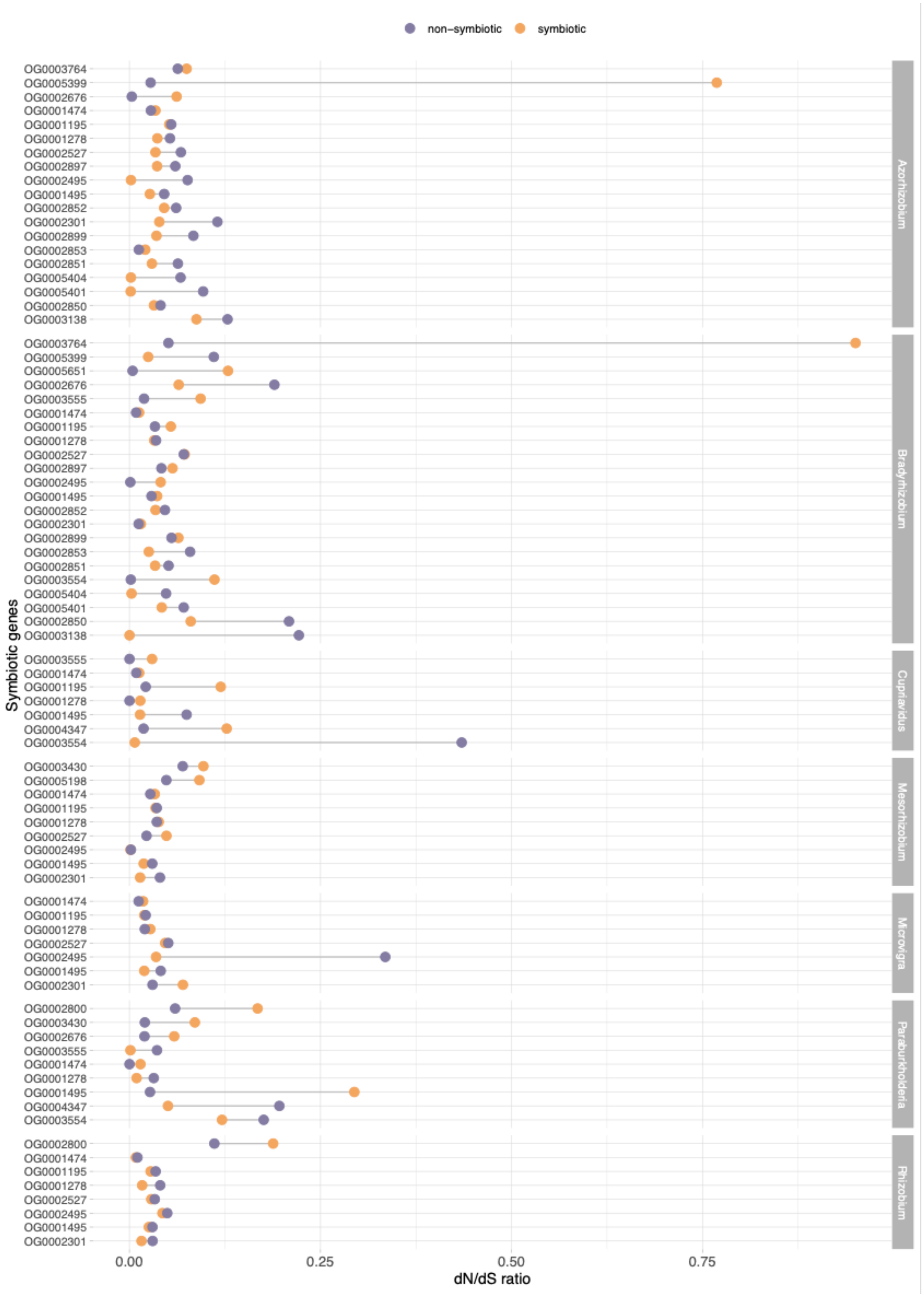


**Supplemental Figure 3. Differences in dN/dS ratios between symbiotic (orange) and non-symbiotic (purple) bacteria species at symbiotic genes.** dN/dS ratios were calculated from free-ratio models in PAML.

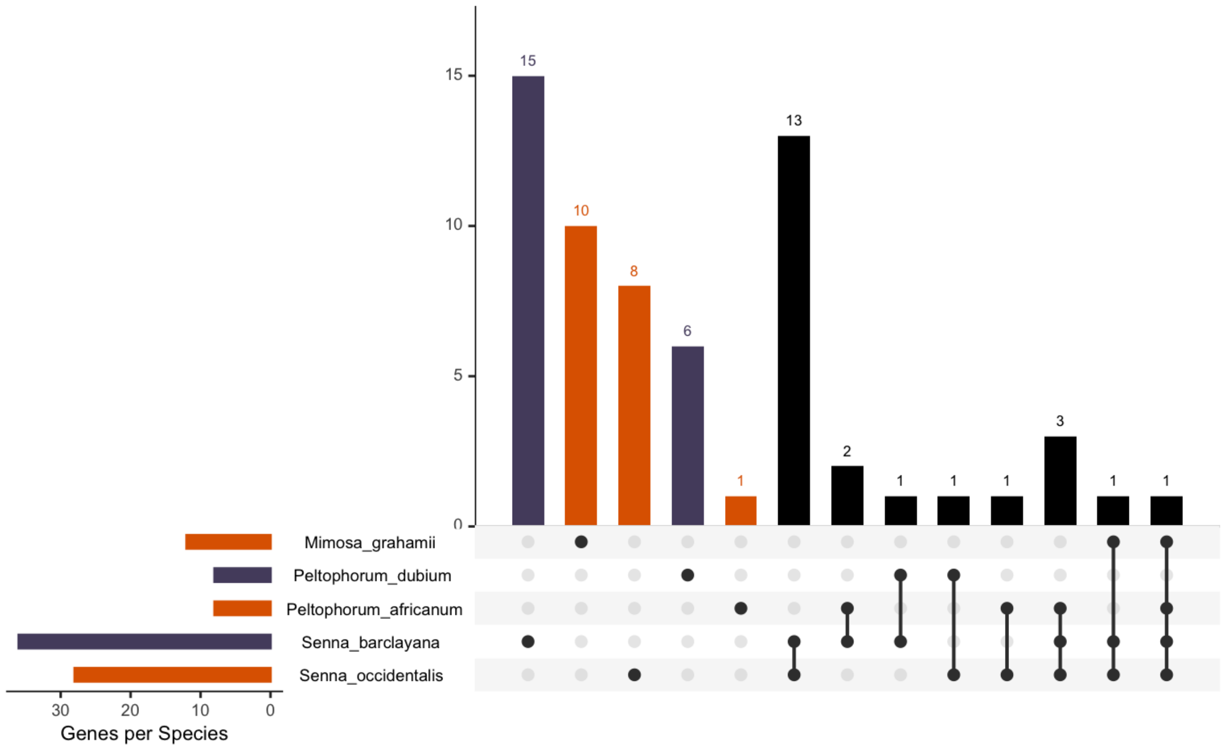


**Supplemental Figure 4. Genes under positive selection in legume genomes.** Mutualistic legumes are in orange and non-symbiotic legume species are in purple.

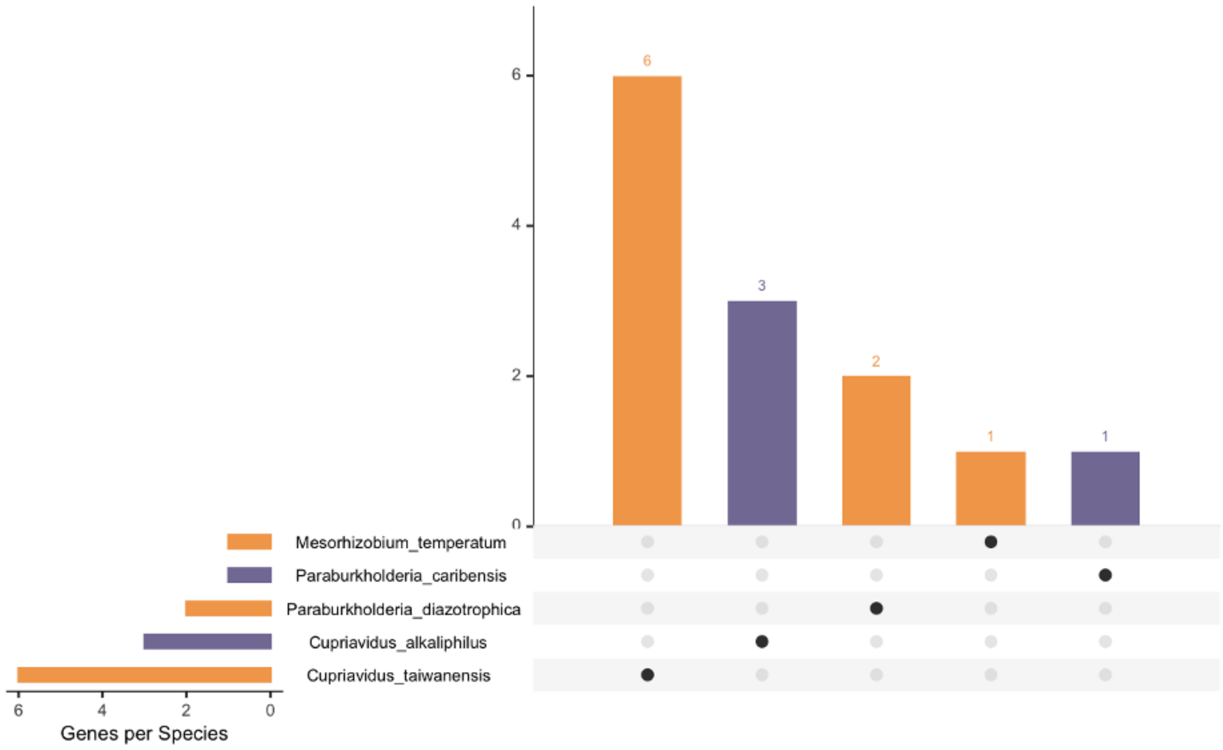


**Supplemental Figure 5. Genes under positive selection in rhizobia genomes.** Symbiotic rhizobia strains are in orange and non-symbiotic species are in purple.
